# Integration and evaluation of magnetic stimulation in physiology setups

**DOI:** 10.1101/2022.07.02.498434

**Authors:** Malte T. Ahlers, Christoph T. Block, Michael Winklhofer, Martin Greschner

**Affiliations:** Department of Neuroscience, Carl von Ossietzky University Oldenburg, Germany; Institute of Biology and Environmental Sciences, Carl von Ossietzky University Oldenburg, Germany

## Abstract

A large number of behavioral experiments have demonstrated the existence of a magnetic sense in many animal species. Further, studies with immediate gene expression markers have identified putative brain regions involved in magnetic information processing. In contrast, very little is known about the physiology of the magnetic sense and how the magnetic field is neuronally encoded. *In vivo* electrophysiological studies reporting neuronal correlates of the magnetic sense either have turned out to be irreproducible for lack of appropriate artifact controls or still await independent replication. Thus far, the research field of magnetoreception has little exploited the power of *ex vivo* physiological studies, which hold great promise for enabling stringent controls. However, tight space constraints in a recording setup and the presence of magnetizable materials in setup components and microscope objectives make it demanding to generate well-defined magnetic stimuli at the location of the biological specimen. Here, we present a solution based on a miniature vector magnetometer, a coil driver, and a calibration routine for the coil system to compensate for magnetic distortions in the setup. The magnetometer fits in common physiology recording chambers and has a sufficiently small spatial integration area to allow for probing spatial inhomogeneities. The coil-driver allows for the generation of defined non-stationary fast changing magnetic stimuli. Our *ex vivo* multielectrode array recordings from avian retinal ganglion cells show that artifacts induced by rapid magnetic stimulus changes can mimic the waveform of biological spikes on single electrodes. However, induction artifacts can be separated clearly from biological responses if the spatio-temporal characteristics of the artifact on multiple electrodes is taken into account. We provide the complete hardware design data and software resources for the integrated magnetic stimulation system.

## Introduction

The Earth’s magnetic field is used across many animal species for navigation, including migratory birds, sea turtles, salmon, lobsters, desert ants, and moths (1–6). Currently, there is increasing interest in the magnetic sense of animals, partly driven by recent advances in understanding the quantum mechanical process likely underlying the remarkable ability of migratory birds to sense the earth’s magnetic field (7). Furthermore, recent studies suggested several candidate brain structures for magnetic cue processing in birds (8–14). Most of these studies, however, do not provide deeper insight into the physiological mechanisms underlying magnetoreception, since they were mainly focused on the expression of corresponding immediate early genes. To date, only a few *in vivo* electrophysiological studies on the magnetic sense are available. Early extracellular recordings that detected magneto-sensitive neurons in the pigeon’s optic tectum (15) turned out to be irreproducible in a technically well controlled replication study (16). Cells in the vestibular brainstem of head-restraint pigeons exposed to sweeping magnetic field stimuli were found to encode magnetic field direction, intensity, and polarity (17). This potentially highly relevant study thus far has been confirmed independently only at the level of immediate early gene expression (13). *In vivo* electrophysiology work detected electrophysiological responses to magnetic fields in the ros V nerve of rainbow trout (18), consistent with abolished conditioned magnetic field responses on trout with anesthetized trigeminal terminals in the snout region (19). However, we are not aware of independent electrophysiological replication attempts.

Thus, *in vitro* and especially *ex vivo* physiological experiments have the potential to close the knowledge gap to the largely unknown underlying cellular and neuronal mechanisms. For these experiments, varying magnetic stimuli have to be presented to the studied specimen while physiological responses (e.g. neuronal activity) are recorded. Here, it is critical to have full control over the magnetic stimuli, i.e., to generate stimuli with the desired properties and to verify that they actually have the desired properties. However, the generation and evaluation of magnetic stimuli for *in vitro* physiology, like single- and multi-electrode extracellular recordings, intracellular recordings, patch-clamp recordings, calcium imaging, and the like, share a number of specific methodological problems. In particular, inherently strong space constraints and the presence of ferromagnetic materials inside the recording setup make it demanding to integrate defined magnetic stimuli.

Common approaches for the generation of spatially homogeneous magnetic stimuli apply coil system designs according to Helmholtz, Lee-Whiting, Merrit, Alldred and Scollar, or Rubens (20). While we concentrate on a square Helmholtz-type coil arrangement here, the findings presented in this paper can be generalized to other systems. In case of Helmholtz coils, for each axis of stimulation, a pair of coils is needed. Hence, full control of the magnetic stimulus in three spatial dimensions requires a total of six coils. This poses a problem for *in vitro* physiology setups, since these are typically space-constricted, in particular, the space around the studied specimen is limited. Often, it is unavoidable to diverge from ideal Helmholtz conditions and place the specimen off-center between the coils, to deviate from the ideal relation between coil distance and coil radius, or to choose a non-ideal coil geometry. These factors potentially degrade the magnetic field homogeneity at the location of the specimen. Furthermore, ferromagnetic components in proximity to the site of stimulation disturb the magnetic field and thereby deteriorate field homogeneity. While it is obviously advisable to reduce ferromagnetic materials inside a magnetic stimulation setup as far as possible, it is rarely possible to avoid them entirely or it is prohibitively expensive. Electronic devices, like preamplifiers or microscope objectives, are situated in close proximity to the site of recording and in most cases contain small amounts of ferromagnetic components. Additionally, electrophysiological setups are mostly installed inside of Faraday cages to shield them from external electromagnetic disturbances. These are often made of ferromagnetic steel due to their better shielding properties for low frequency electromagnetic fields in comparison to non-ferromagnetic materials. Also, the building the experiments are performed in, might contain ferromagnetic structural elements. The presence of such materials distorts the magnetic field inside the coils and potentially degrades the stimulus quality. The stimulus properties at the location of the specimen thus have to be verified. However, also in this regard, the spatial constraints of *in vitro* physiology setups are problematic. The specimens are typically relatively small (in the order of several millimeters). Most commercially available magnetometers (e.g. fluxgate magnetometers) are larger than typical physiological recording chambers, more so if three axes of simultaneous measurement are needed for full spatial characterization of magnetic fields. Smaller devices (e.g. Hall sensors) are typically not sensitive enough. Hence, these large magnetometers do not fit into the site of recording, at least not without modifying or removing setup components. It is, however, important that all setup parts are in the same configuration as during the experiment, since they potentially alter the magnetic field. Moreover, the characterization of field homogeneity across the specimen is limited with sensors whose spatial integration area is larger than the specimen itself. In addition, in multi-axis fluxgate magnetometers, the axes of measurement are often significantly offset from each other (tens of millimeters) due to the size of the sensory structures.

Here, we present a three-axis magnetometer design based on anisotropic magnetoresistive (AMR) sensors commonly used in smartphones. Since these are used as compass sensors, they are capable of measuring magnetic fields with a sensitivity of fractions (typically in the 100th to 1000th) of the earth’s magnetic field in three axes. They are small enough to fit in common recording chambers of setups for physiological research and have small spatial integration areas, making them well suited for the purpose. Moreover, we present a design of a coil-driver circuit that is able to flexibly generate magnetic stimuli, enabling analysis of biological responses to non-stationary fast changing stimuli. We provide the complete design data for both devices, i.e. layouts of the printed-circuit-boards, microcontroller firmware, and high-level calibration software (www.github.com/mtahlers/magstim). The Helmholtz coil driver module can be built by a person with entry-level practical electronics skills. Building the magnetometer module is slightly more demanding due to the smaller size of the used components. We present a calibration routine for the magnetometer and the Helmholtz coil system that compensates for stationary soft- and hard-iron distortions. Finally, we demonstrate the effect of magnetic induction artifacts in relation to electrophysiological recordings from retinal ganglion cells and furthermore demonstrate the separability of neuronal spikes and magnetic stimulation artifacts.

## Materials and methods

### Miniature vector magnetometer

The presented vector magnetometer is based on the QMC5883L 3-axis magnetic sensor (21). Its package measures 3 mm x 3 mm x 0.9 mm. The sensor deploys the anisotropic magnetoresistance (AMR) principle, i.e. the change of electrical resistance of a nickel-iron (permalloy) thin-film element under the influence of an externally applied magnetic field. Four magnetoresistive elements are connected as a Wheatstone bridge, forming one axis of sensitivity (22). Hence, three of these structures are arranged perpendicularly to form a full vector magnetometer. In contrast to older AMR sensors, the QMC5883L offers an integrated analog front-end and a 16-bit analog-to-digital converter so that the measured magnetic data can be digitally accessed via an I^2^C bus by a small microcontroller. This significantly reduces design demands and increases robustness since no external analog circuitry has to be developed. The QMC5883L applied here is a derivative of the HMC5883L by Honeywell (23), using the same magnetoresistive technology but with its digital resolution increased from 12 to 16 bit. While the pin-out and the required external circuitry of both sensors are the same, the internal programming register structure is different, making the sensors not a direct replacement on the firmware level.

The QMC5883L is commonly available soldered on break-out printed circuit boards (PCBs) giving easy access to its electrical connections and providing the circuitry for basic operation. However, we designed our own carrier PCB for the sensor for two reasons: Firstly, we wanted to minimize the size of the sensor unit as much as possible. Secondly, we found some of the electronic components on the commercial break-out PCBs to be strongly ferromagnetic and therefore to disturb the magnetic measurement in an easily avoidable manner. This was the case for the pins of some of the ceramic capacitors and for the connection pin header. We chose a two-part construction for the sensor unit (Fig. 2A). A small PCB only carrying the QMC5883L was soldered perpendicular to an adapter PCB that is populated with the required decoupling capacitors in close proximity to the sensor. It also provides the contact points for connecting the data cable to the interface unit. By this means, a sensor unit with a diameter of 6 mm was constructed. We provide the detailed schematic and a printed circuit board layout (www.github.com/mtahlers/magstim).

The sensor data is transmitted via the synchronous serial bus I^2^C. We chose the 8-bit AVR microcontroller ATmega168 (24) to control and read out the QMC5883L. The data output rate, the oversampling factor, and the sensitivity of the measurement are adjusted through a control register of the sensor. In this register, we set the sensor to the highest data output rate, 200 Hz, to the lowest oversampling, 64x, and to the highest sensitivity of the measurement, ±2G (see also results).

The UART-to-USB bridge IC FT232R (25) is used in our module to connect the AVR microcontroller to a standard PC. By this means, the connection to the magnetometer integrates as a virtual serial port into the operating system of the used host computer.

For the sake of flexibility, the microcontroller transmits the QMC5883L data to the USB-attached high-level computer without any data processing. All further conditioning, e.g. averaging and filtering, of the gathered magnetic data is thus performed on the host PC. A simple data protocol was chosen for data transmission. Since the sensor outputs the measured values as a 16-bit signed integer data type, a space-separated triplet of ASCII encoded numbers ranging from -32768 to 32767 is transmitted, with each number representing the value of one of the axes. Each data triplet is terminated by the ASCII characters <CR> and <LF>.

The magnetometer connection integrates as a generic COM port into the used operating system. Hence, practically any operating system and programming language can be used to receive the data. We used Matlab (Mathworks, MA, USA) to receive and process the magnetometer measurement data.

### Calibration of the vector magnetometer

Ideally, a vector magnetometer measures the magnetic field projections (*B*_*x*_, *B*_*y*_, *B*_*z*_) onto three independent orthogonal spatial axes (*x, y, z*) with equal gain and zero offset. Hence, if an ideal magnetometer is arbitrarily rotated in a spatially uniform magnetic field (as the ambient Earth’s magnetic field), all measured coordinates lie on a sphere centered at *B*_*x*_*=B*_*y*_*=B*_*z*_*=0* and with a radius equal to the total field intensity, sqrt (*B*_x_^2^+*B*_y_^2^+*B*_z_^2^). However, in practice, different factors contribute to non-ideal behavior of magnetometers. In AMR sensors, each axis is measured by four magneto-resistive elements arranged as a Wheatstone bridge. Typically, these elements vary slightly in resistance which leads to a zero-field offset voltage of the bridge. The offset does not change value or polarity if the magnetic stimulus varies and can be assumed to be constant over the lifespan of the sensor (26), however, it varies between different sensors. The offsets of all three axes can be viewed as an addition of a constant error vector to the measurement. Furthermore, the gain might differ between the different axes of a sensor. This leads to a deformation of the ideal measurement sphere into an ellipsoid. In principle, a sensor axis can be weakly sensitive to magnetic fields orthogonal to its main axis, i.e. showing so-called cross-axis sensitivity (27). This effect can result from uneven magnetization that the permalloy in AMR sensors might acquire over time. However, as most modern integrated AMR magnetometers, the QMC5883L has a built-in degaussing functionality that resets any magnetization bias by applying a magnetic reset-pulse to the permalloy strips prior to each measurement. In addition to these sensor-intrinsic error sources, extrinsic factors contribute to the perturbation of the measured magnetic field. Any static magnetic field source fixed to the reference frame of the magnetometer will bias the measured field, e.g. magnets or pieces of magnetized material on the sensor’s circuit board. These so-called hard-iron distortions can be described as an offset and together with the zero-field offset mentioned above can be combined to give a single offset vector for the calibration procedure. In addition, so-called soft-iron distortions are caused by the induction of magnetic fields into normally unmagnetized ferromagnetic objects in proximity to the sensor, distorting the magnetic field lines. The induced soft-iron fields are proportional to the relative magnetic permeability of the material, thus especially iron and nickel on the circuit board tend to induce this effect. If these materials are present in proximity to the sensor, their effect is to stretch and tilt the sphere of ideal measurements.

If an uncalibrated magnetic sensor is arbitrarily rotated in a spatially uniform magnetic field, the sensor’s raw x-, y-, and z-component data describe a somewhat distorted sphere, i.e. an ellipsoid with its center slightly offset. The parameters describing this ellipsoid can be derived analytically (28). Here, a vector ***V***_bias_represents the offset of the sphere (bridge offsets and hard-iron effects). The effects of soft-magnetic distortions, different gain along each axis, and potential uncompensated cross-axis sensitivity, can all be combined into a single 3 × 3 matrix **W**, so that the uncalibrated readings ***V***_raw_of the magnetic field (here the raw digital output value triplets of the sensor) can be mathematically represented as

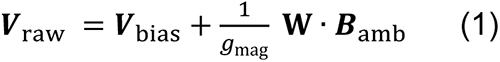

where ***B***_amb_ is the ambient magnetic field (in units of flux density, T), which can be retrieved by undoing the effects of ***V***_bias_and **W**, i.e.,

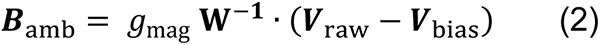

where **W**^−**1**^ is the inverse of **W** and *g*_mag_ (in units of flux density per least significant bit of sensor output, T/LSB) is the scale factor to transform the read-out sensor values into absolute values of ***B***_amb_on the basis of a reference measurement with a calibrated magnetometer.

Matlab’s Sensor Fusion and Tracking Toolbox (Mathworks, MA, USA) provides the *magcal* function. It determines the calibration parameters corresponding to **W**^−**1**^ and ***V***_bias_for sensor raw data that resulted from freely rotating a magnetometer sensor in a homogeneous magnetic field. Alternatively, other functions based on ellipsoid fitting can be used, e.g., the code provided with the application note “Ellipsoid or sphere fitting for sensor calibration” (29).

The earth’s magnetic field strength at the location of the calibration data acquisition was measured with a calibrated commercial magnetometer (FVM400, Macintyre Electronic Design Associates).

### Helmholtz Coils

Helmholtz coils are commonly used to generate nearly uniform magnetic fields in the central region of the coil system. Each pair of coils in a tri-axial coil system ideally consists of two parallel, equally sized circular or square coils with an identical number of windings. When a current flows through the windings, the magnetic fields of both coils combine. If the distance between two circular coils of a pair is equal to their radius *r*, the resulting field has a high homogeneity in the middle between the coils. For two circular coils of radius *r*, distance *d* = *r, n* windings, powered by current *I*, and *µ*_*0*_ being the vacuum permeability, the magnetic field strength *B* at the midpoint between the coils is:

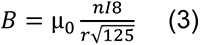

For quadratic coil pairs of side length *a*, best field homogeneity is achieved when choosing the separation distance to be *d* ≈ 0.5452 *a* (Kirschvink, 1992). The quadratic x-, y-, and z-Helmholtz coil pairs, purpose-built for our exemplary setup, had a side length of *a*_x_ = 223 mm, *a*_y_ = 400 mm, and *a*_z_ = 162 mm. For each pair of coils, the distances *d* were chosen to satisfy the condition *d*/*a* ≈ 0.5452 for maximum field homogeneity. To allow for field blanking (see next paragraph), the Helmholtz coils in our exemplary setup were double wound. Turns per coil winding were *N*_x_ = 32, *N*_y_ = 53, *N*_z_ = 19. DC resistances of the coil pairs were *R*_x_ = 10.1 Ω, *R*_y_ = 29.9 Ω, and *R*_z_ = 4.4 Ω, inductances were *L*_x_ = 1.05 mH, *L*_y_ = 5.16 mH, and *L*_z_ = 0.24 mH. Resistances and inductances were measured by a Fluke PM6306 LCR-meter (Fluke, WA, USA).

Time-dependent magnetic stimuli are prone to produce induction artifacts in electronic devices. This applies all the more to electrophysiological experiments, which rely on high impedance voltage measurements. The research field of magnetobiology has suffered several drawbacks, some of which had resulted from improper controls for magnetic field exposure conditions (30). A recommended control is referred to as sham exposure, which consists in blanking the magnetic field of an electrically activated coil pair. To allow for blanking, two independent wires are wound in parallel onto the coil spools that can be connected serially or anti-serially during operation. This coil type is commonly referred to as double-wrapped (20). When connecting both windings in series, their magnetic fields constructively add up. By connecting them anti-serially, the magnetic fields produced around the wires cancel out. For a sham exposure the identical electrical power is applied to the coil as in the real magnetic field exposure. Artifacts induced by electrical activation of the coils can be identified by this means. However, artifacts induced by the magnetic field in the real exposure condition can obviously not be addressed by this control and need to be identified in the context of the experiment (see “Induction Artifacts” in Results section).

Together with the magnetic field an electrical field is produced by the windings of the Helmholtz coils. However, due to the geometry of the coils the resulting electric field is small in the center between the coils. In contrast to coils, parallel plates are typically used to produce homogenous electrical fields. As is the case for the magnetic field, the electric field is canceled out in the sham condition due to the opposite current polarity in neighboring coil windings.

### Helmholtz Coil Drivers

The magnetic field strength produced by Helmholtz coils is proportionally dependent on the current flowing through its windings. Ohmic losses in the powered coil windings increase the temperature of the coil which in turn increases the resistance. If the coil is powered by a constant voltage, the temperature driven rise in resistance will decrease the coil current and thus the magnitude of the generated magnetic field over time. It is therefore advisable to power the coils by a constant current source. Any change in the resistance of the coil due to temperature variation will thereby be counteracted by voltage adjustments by the source, keeping the current and thus the magnetic field strength at the desired level.

Figure 1B shows the structure of the voltage-controlled current source implemented for the Helmholtz coil driver described here. The operational amplifier controls the shunt voltage at the resistor *R*_Shunt_ to be equal to the level at its positive input *U*_control_ by adjusting the output voltage *U*_out_ accordingly. According to Kirchhoff’s first law, the current through *R*_Shunt_ is the same as through the Helmholtz coil winding (neglecting very small input currents into the operational amplifier due to its high input impedance). The coil current *I*_coil_ is thus

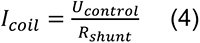

Hence, ***I***_*coil*_ can be proportionally controlled by ***U***_*control*_ with the proportionality factor defined by 1/***R***_*shunt*_. Any variation of the control voltage will directly be translated into variations of the coil current and thus the magnetic field strength.

We provide the detailed schematic of the module and a printed circuit board layout (www.github.com/mtahlers/magstim). The design only utilizes trough-hole-technology components, making it easy to assemble also for an electronics amateur.

For the sake of simplicity and robustness of the control circuitry, we chose the OPA548 power operational amplifier (31). While most operational amplifiers can deliver only small output currents, the OPA548 provides an integrated power output stage capable of sourcing and sinking up to ±3 A continuously. This reduces the design effort and component count of the module since no additional discrete power output stage is needed. The current range of ±3 A is well suited for driving medium sized coils as typically needed for physiological experiments.

Any stabilized bipolar DC voltage source of sufficient voltage and current output can be used to supply the driver module through its “POWER” connector. The lowest required voltage for the driver module to operate can be estimated on the basis of the required DC coil voltage and the dropout voltage of the operational amplifier. For example, if a maximum current of ±1 A has to flow through a 5 Ω coil pair, a voltage of ***U*** = ***R*** · ***I*** = ±5*V* would be needed according to Ohm’s law. The operational amplifier has a maximum dropout voltage of approx. 4 V (31), it can thus output roughly 4 V less than its supply voltage. For this example, the module should be supplied with at least ±9 V, accordingly. However, step-like changes of the coil current require larger voltages since *di/dt* = *U/L*, i.e., the larger the supply voltage, the faster is the transition time between different coil currents and thus magnetic field strengths. On the other hand, if a steady state current is reached after a step, a power proportional to the difference of the supply voltage and the actual coil voltage is thermally dissipated by the operational amplifier (*P*_*diss*_ = Δ***U*** · ***I***). Practically, a larger difference between the supply voltage and the needed DC coil voltage will lead to more heat production by the amplifier. In any case, a properly dimensioned heat sink should be attached to the OPA548, and in some cases, active cooling by a fan might be necessary. The chosen current shunt resistor exhibits a very small temperature coefficient, attaching a heat sink further improves the overall thermal stability. Moreover, it is advisable to let the amplifier thermally settle after power-on for some time. For the presented setup, a field step of 50 µT after being powered off for 2.5 hours at room temperature resulted in an error magnitude of 0.11 ± 0.06 µT for the first second. A two-hour random stimulation of 2 second segments of uniformly distributed field vectors of 50 µT, similar to that in Fig. 5 and 6, resulted in a mean error magnitude of 0.09 ± 0.04 µT. This fluctuation is in the same range as the background field stability measured with the magnetometer in the building.

The Helmholtz coil pair has to be connected to the “COIL” output of the driver module. For field blanking, the amplifier circuitry allows to reverse the polarity of one coil winding by means of a relay. The first winding has to be connected between connection points 1A and 1B of the circuitry and the second winding between 2A and 2B. These signals are provided by the 9-pin D-sub connector on the driver board. The onboard relay can then reverse the polarity of winding B. The “COIL POL.” input controls the polarity reversal relay for the half-windings. The relay is powered from the positive coil supply voltage V+ through a linear voltage regulator. If just a single continuous coil winding is used, 2A should be shorted to 2B by a low resistance connection, e.g., a short wire of sufficient diameter.

The shunt voltage, i.e., the voltage directly proportional to the current through the coils, is provided at the “U_SHUNT” output of the module. It can be used to indirectly monitor the coil current. However, since it is not buffered on the module, high impedance voltmeters should be connected to it. The current measurements presented in Figure 3A and D were obtained by recording the voltage on the “U_SHUNT” output.

The voltage controlling the magnetic field strength has to be connected to the “FIELD CNTRL” BNC input of the driver module. This voltage is divided by the potentiometer R4 before reaching the positive input of the operational amplifier, allowing to adjust the sensitivity of the module. We used three digital-to-analog output channels of a NI-USB 6343 interface (National Instruments, TX, USA) to generate the control voltages, connected to the x-, y-, and z-coil amplifier’s “FIELD CNTRL” input, respectively. The fourth analog-out channel of the interface was used to provide a trigger signal. Three digital output channels of the interface were connected to the “COIL POL.” inputs to control field blanking. The four analog output channels of the NI-USB 6343 provide 16-bit resolution at a maximum data output rate of 719 kSample/s per channel.

The frequency compensation of the amplifier circuitry is adjustable by the trimmer potentiometer R7. For adjustment, a slow (e.g. 100 Hz) square wave control voltage can be connected to the “FIELD CNTRL” BNC connector of the driver. The resulting time course of the coil current can then be monitored by means of the voltage at the U_SHUNT connector. Depending on the inductance of the attached Helmholtz coil, a low compensation might lead to a strong overshoot and ripple of the coil current in response to a step change, while a high compensation slows down the response, i.e., making it less steep. A compromise between a fast but potentially overshooting response and a non-overshooting but sluggish step response has to be found, depending on the application (Fig. 3B, C). In any case, it is very unlikely to directly elicit any spiking activity by magnetic stimulation with our system as in transcranial magnetic stimulation. The magnetic field’s maximum rate of change attainable with our coil driver is approx. 5000 times smaller than those applied in typical transcranial magnetic stimulation systems, and the typically applied absolute field magnitudes are 60000 times smaller (e.g. (32)).

### Calibration of the coil system

An animal moving freely in a spatially uniform ambient magnetic field experiences directional changes of the field vector. To mimic this inside a research setup, a magnetic vector of a fixed magnitude with different spatial orientations needs to be produced. However, the magnetic field generated by Helmholtz coils is subject to the same distortional effects as described for the magnetometer calibration. Here, hard- and soft-iron-distortion by magnetized or ferromagnetic setup components lead to similar deformational effects. Thus, as with the magnetometer calibration, the goal is to invert these deformations. In this case, however, solving for the inversion parameters can be simplified since the corresponding control voltage vector and resulting magnetic field vector pairs are known.

The effective magnetic field, ***B***_eff_(***r***) measured with the calibrated magnetometer at any point *r* in the setup can be represented as a vector sum of the following contributions,

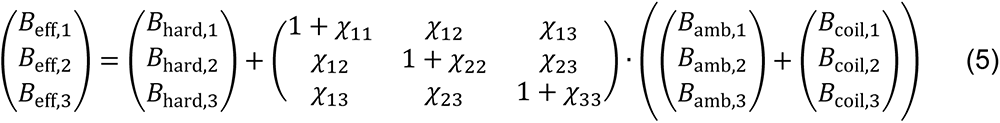

Where ***B***_hard_(*r*) is the non-uniform stray field due to the often unavoidable presence of hard-magnetic components in and around the setup, whose magnetization is invariant of the applied magnetic field. While ***B***_hard_(*r*) typically has only a few small localized sources, both the ambient magnetic field, ***B***_amb_, and coil field, ***B***_coil_, act on a much larger scale and thus can efficiently magnetize soft-magnetic material, e.g., the electric shield around the setup. The magnetization induced in the soft-magnetic material by ambient and coil field may diminish, reinforce, or deflect the magnetic field measured at ***r***. The anisotropy terms χ_*ij*_(*i* ≠ *j*) in Eq. (5) account for deflection, while the isotropic terms χ_*ii*_ describe a reinforcement (χ_*ii*_ > 0) or diminishment (χ_*ii*_ < 0) of the field that would be present without soft-magnetic material. We assume the induced magnetization to be linear because most soft magnetic materials behave linearly in the magnetic field range of interest here (*B* < 200 μT).

The relationship between the voltage applied to the coil and the generated field can be expressed most generally as 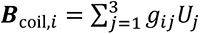 where ***U***_*j*_ is the control voltage applied to the driver of the *j*-th coil axis and *g*_*ij*_ is the voltage-to-field conversion matrix for the coil system (units: T/V). Even in an ideal coil system, consisting of three pairs of identical coils, whose symmetry axes are orthogonal to one another and intersect in a single point, *g*_*ij*_ would be a diagonal matrix only in the center. In all other cases, the off-diagonal elements are finite.

Given a spatially uniform ambient field, there are still 15 unknowns (***B***_hard,i_, χ_*ij*_, *g*_*ij*_) at any point in the setup, but these need not be determined explicitly if the sole task is to find the triplet of control voltages (***U***_1_, ***U***_2_, ***U***_3_) for the coil drivers in order to generate a defined field vector ***B***_eff_ at the position of the specimen. In this case, Eq. (5) can be rewritten as a single matrix multiplication,

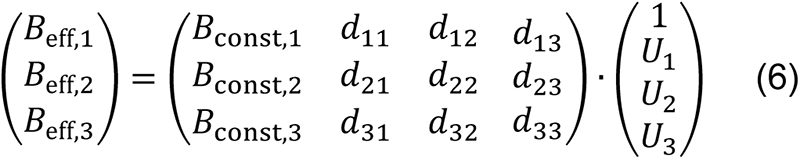

where 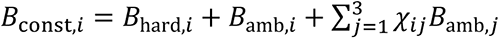 are the constant terms at the given position, which are directly obtained when measuring the effective field with the coils turned off before the calibration, i.e., *B*_const,*i*_ = *B*_eff,*i*_(***U***_1_ = ***U***_2_ = ***U***_3_ = 0). Therefore, we have

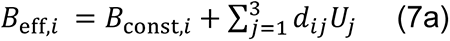

or in matrix notation,

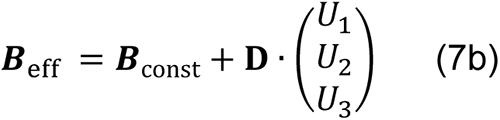

The task during calibration is to determine the numerical values of the elements *d*_*ij*_ of matrix **D**. In the calibration routine, a multitude of combinations of voltage triplets (dependent variables) are applied while taking the respective field readings (response variables), i.e.,

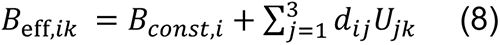

where the index *k* to *B*_eff,*ik*_ and ***U***_*jk*_ refers to the *k*-th triplet of voltages. The *d*_*ij*_ values are then determined by fitting each Cartesian component of Eq. (8) to the data, using linear regression. The matrix **D** then is inverted to obtain the voltage triplets needed to set the field ***B***_eff_, i.e.,

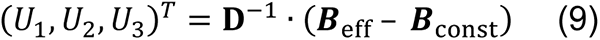

The mathematical similarity between Eq. (2) for the magnetometer calibration and Eq. (9) for the coil calibration reflects the conceptual similarities between the derivations. Therefore, to visualize Eq. (8), we follow a similar approach as in the magnetometer calibration, now applying a large number of different control voltage triplets of constant magnitude 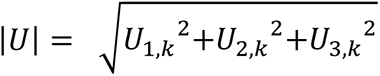, so that **D** acts on a spherical surface in a three-dimensional “voltage space” centered at the origin. The resulting distribution of ***B***_eff_ is shifted from the origin by the offset field ***B***_const_ and is typically deformed into an ellipsoid (effect of **D**). After successful calibration, the values of (***U***_1_, ***U***_2_, ***U***_3_) can be set such that the resulting distribution of ***B***_eff_ is spherical and centered at the origin so that one can achieve different effective field directions at the location of the sample while keeping the effective magnetic field intensity constant.

### Multielectrode recordings with simultaneous visual and magnetic stimulation

We extracellularly recorded electrical activity from retinal ganglion cells of the common quail (*Coturnix coturnix*) under simultaneous visual and magnetic stimulation. All experiments were performed in accordance with the institutional guidelines for animal welfare and the laws on animal experimentation issued by the European Union and the German government. Segments of pigment epithelium attached retina were placed flat, ganglion cell side down, on a planar array of extracellular microelectrodes. The electrode array consisted of 512 electrodes and covered a rectangular region of 1890 µm x 900 µm (35). The retina was submerged in Ringer’s solution (100 mM NaCl, 6 mM KCl, 1 mM CaCl_2_, 2 mM MgSO_4_, 1 mM NaH_2_PO_4_, 30 mM NaHCO_3_, 50 mM Glucose), bubbled with carbogen (95% O_2_ and 5% CO_2_), pH 7.5 (33). Recordings were analyzed offline to isolate the spikes of different cells, as described previously (34). Briefly, candidate spike events were detected using a threshold on each electrode, and the voltage waveforms on the electrode and neighboring electrodes around the time of the spike were extracted. Clusters of similar spike waveforms were identified as candidate neurons if they exhibited a refractory period. Duplicate recordings of the same cell were identified by temporal cross-correlation and removed.

Magnetic stimulation consisted of a random sequence of rapid switches between 12 evenly distributed magnetic field vectors of a magnitude of 50 µT (Fig. 6D, insert). A new magnetic field vector was chosen every 200 ms. A visual noise stimulus, updating randomly and independently over time at 120 Hz, was used to calculate the spike-triggered average stimulus and to characterize the response properties of the recorded cells (Fig. 6F).

The electrical image of a ganglion cell is the average spatiotemporal spike waveform recorded across the electrode array during a spike (35) (Fig. 6 A-E). The electrical image of the induction artifact was calculated as the average waveform across the electrode array during a switch of the magnetic stimulus. The electrical images of the induction artifact were separated for all 12*12 stimulus transitions (Fig. 6D). In the presented setup, field changes in x- and y-coil direction produced the strongest induction. No influence of the individual routing of the electrode traces on the array was apparent. A linear fit of the peak amplitude to the magnetic field transition was used as a color map. Recordings were bandpass filtered (80 Hz - 2 kHz) prior to averaging.

Unless stated otherwise, all measurements are reported as the mean ± standard deviation.

## Results

Often, the space-restricted nature of typical physiology setups hinders the integration of the field-generating electromagnetic coil system. Therefore, ideal placement of the coil system is rarely possible. Figure 1A shows a typical setup for extracellular multielectrode recording from the retina with an integrated three-axis magnetic stimulation system. During recording, the specimen is situated at the position indicated by the letter S. Ideally, the surrounding three pairs of Helmholtz coils would be centered around this location. However, the presence of the recording electronics, microscope, and stimulus projection optics made it necessary to deviate from this situation, i.e., to place the coils off-center from the specimen, while maintaining the optimal radius-to-distance ratio (see methods “Helmholtz coils”). Additionally, small amounts of ferromagnetic materials are present in the nearby setup components, potentially disturbing the homogeneity of the produced magnetic field. Under these non-ideal conditions, it is particularly important to evaluate the magnetic field magnitude and homogeneity, since they might be compromised by these design prerequisites. The region of interest for the magnetic measurement is typically just a few millimeters in size, and the surrounding recording chambers are typically just several tens of millimeters wide. Therefore, a sufficiently small magnetometer is necessary to characterize the magnetic field at the position of the preparation without removing components of the setup. Due to their size, commercial magnetometers like fluxgate magnetometers do not fit into recording chambers and integrate over too large areas.

**Figure 1:**
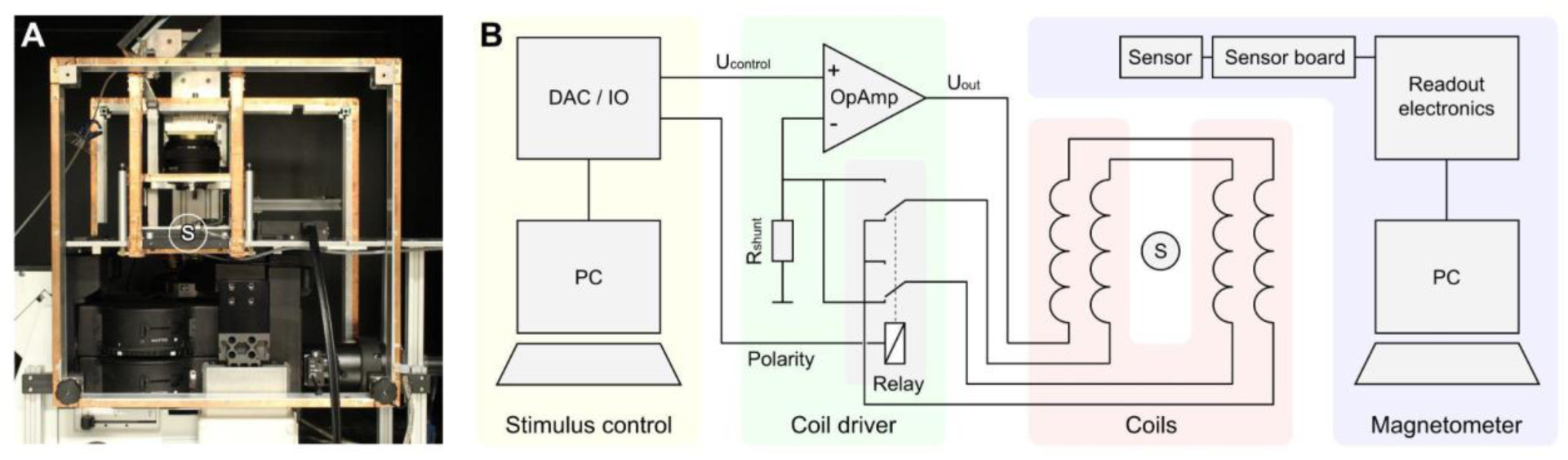
Electrophysiological setup with an integrated three-axis Helmholtz coil system. **A:** Extracellular multielectrode recording setup with integrated coils. **B:** Schematic overview of the setup components for the generation and measurement of magnetic field stimuli. Coil driver and coils are shown for one of three axes. The placement of the specimen is indicated by the letter S in A and B.

### Magnetometer

We developed a magnetometer based on the 3-axis magnetic sensor QMC5883L (21). To benefit from the sensor IC’s small package size of 3 mm * 3 mm * 0.9 mm, the carrier PCB should be as small as possible. Since the device needs decoupling capacitors in close proximity, we chose a two-part construction to obtain a compact design: The sensor is placed on a PCB with a diameter of 6 mm to which a connector PCB is soldered perpendicularly (Fig. 2A). The latter carries the decoupling capacitors and the solder points for the sensor cable. Thereby, a very compact design of the sensor board was achieved, that can be placed in typical recording chambers of physiological setups. The sensor raw data is interfaced to a USB port of a standard PC by readout electronics consisting of a microcontroller and UART-USB bridge IC.

**Figure 2:**
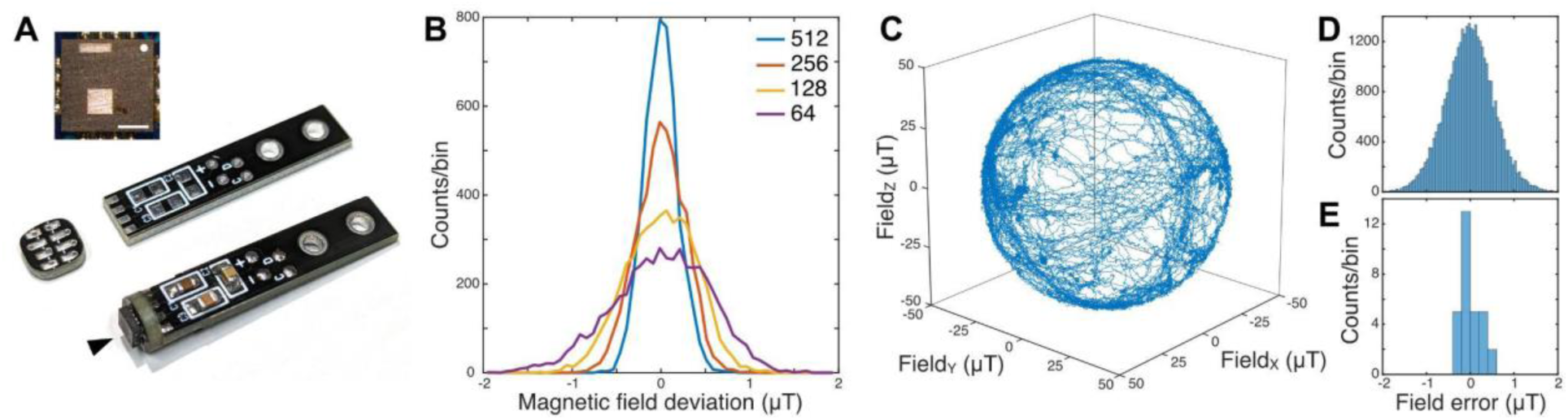
Miniature vector magnetometer. **A:** Two-part sensor board design. The QMC5883L magnetic sensor is marked by an arrowhead. Sensor 3 mm x 3 mm. Inset top left: Position of the sensor dies inside the QMC5883L package, top view onto the package. White dot marks pin 1. Inset scale bar = 1 mm **B**. Noise floor of an exemplary QMC5883L for the four available oversampling settings 64x - 512x. **C**. Trajectory of rotating the sensor in the natural earth magnetic field (49 µT) for approximately 3 minutes after calibration on the smallest oversampling setting. **D**. Residuals between the trajectory data in C and a homogeneous natural field. **E:** Residuals as in D averaged over 15 s to reduce noise for 30 stationary orientations in the natural magnetic field.

We decapsulated a QMC5883L by grinding open its package from the top side, revealing two spatially separated sensor dies, one integrating the x- and y-axis of measurement, one implementing the z-axis of measurement (Fig. 2A inset top left). Since the x-y-structure is monolithically implemented in one die, orthogonality of the x- and y-axis of measurement can be expected to be very good, also before calibration. Non-orthogonality, potentially introduced by imprecise alignment of the two dies, is compensated by the calibration procedure (see methods). The spatial integration area of the sensor structures is significantly smaller than the sensor’s package size, namely roughly one-third of it. Obviously, the overall dimension, as well as the offset between axes, is very small in comparison to fluxgate magnetometers.

We compared four exemplary sensors by applying the calibration procedure as described below to obtain the axes’ gain and offset values of each device. The sensors had gains of 7.85 ± 0.19 nT per least significant bit (n=4 sensors). On average, the magnitude of the offset vector of the four sensors was 1.09 ± 0.49 µT. We determined the noise floor of the calibrated sensors by exposing them stationarily to the earth’s magnetic field (49.0 µT, natural field in Oldenburg, Germany) (Fig. 2 B). The QMC5883L provides internal data averaging over a window of 64, 128, 256, or 512 data points, and data output rates of 10, 50, 100, and 200 Hz, both adjustable via the sensor’s control register (21). While the different data output rates did not have any influence on the noise of the sensor data, larger averaging windows decreased the noise floor, as expected (std 0.54, 0.39, 0.27, and 0.19 µT for 64x, 128x, 256x, and 512x oversampling). All values are in good accordance with the respective datasheet values (21). In our magnetometer firmware, we set the sensor to the fastest output rate (200 Hz) and the lowest oversampling value (64x). By this means, the magnetometer provides the fastest possible response to field changes. To increase the sensor’s precision, averaging over constant field conditions or filtering techniques can be applied downstream in real-time or offline (e.g. Fig. 2E).

### Magnetometer calibration

Optimally, the calibration of the magnetometer is performed in a calibrated coil system that is able to provide arbitrary magnetic fields. If a calibrated coil system is not available, a procedure can be applied that is similar to the calibration routine for smartphone-integrated compasses. Correspondingly, the sensor board of our magnetometer was randomly rotated in the undisturbed earth magnetic field, i.e. sufficiently far away from buildings and other structures potentially containing metal. The magnetic field strength at the location of data collection was 49.0 µT. The x-, y-, and z-axis raw data of the sensor lie on a deformed sphere offset from the origin (see methods). The offset values and deformation matrix were extracted by the calibration routine (see methods). Applying those to the raw sensor output data made them spherical and zero-centered (Fig. 2C). After calibrating the exemplary sensor in the local earth magnetic field (49 µT), the mean field error was close to zero (0.02 µT) and the standard deviation was 0.55 µT (Fig. 2D). To estimate the accuracy of the calibrated magnetometer in the absence of the sensor’s noise floor, we placed the calibrated magnetometer stationarily in multiple arbitrary orientations in the natural magnetic field and averaged data in every position. For the exemplary sensor and calibration procedure, the residuals between the averaged data and a homogeneous natural field were close to zero (0.01 µT) and had a standard deviation of 0.23 µT, which corresponds to an error of 0.47% of the measured magnitude of 49 µT (Fig. 2E).

### Helmholtz Coil Drivers

Helmholtz coils are preferably driven by a constant current source since their generated magnetic field is directly dependent on the current. By this means, changes in the coil resistance, resulting from temperature changes of the coil, do not influence the magnetic output of the coils. The core of the coil driver design is an operational amplifier with an integrated power output stage (Fig. 1B). The amplifier circuitry translates a control voltage into a proportional current, that is powering the Helmholtz coils. The proportionality factor of the voltage-to-current conversion is defined by a shunt resistor.

To characterize the linearity of the coil amplifier, we increased the control voltage from a value resulting in a magnetic field strength of approx -100 µT to a value resulting in approx. 100 µT. We measured the voltage drop at the current shunt to obtain the corresponding coil current (*I*_*coil*_ *= U*_*Shunt*_ */ R*_*Shunt*_). Simultaneously, we measured the generated magnetic field magnitude by the analog output of a commercial magnetometer (FVM400, Macintyre Electronic Design Associates). The deviation from a linear fit to the data was minimal (Fig. 3A).

**Figure 3:**
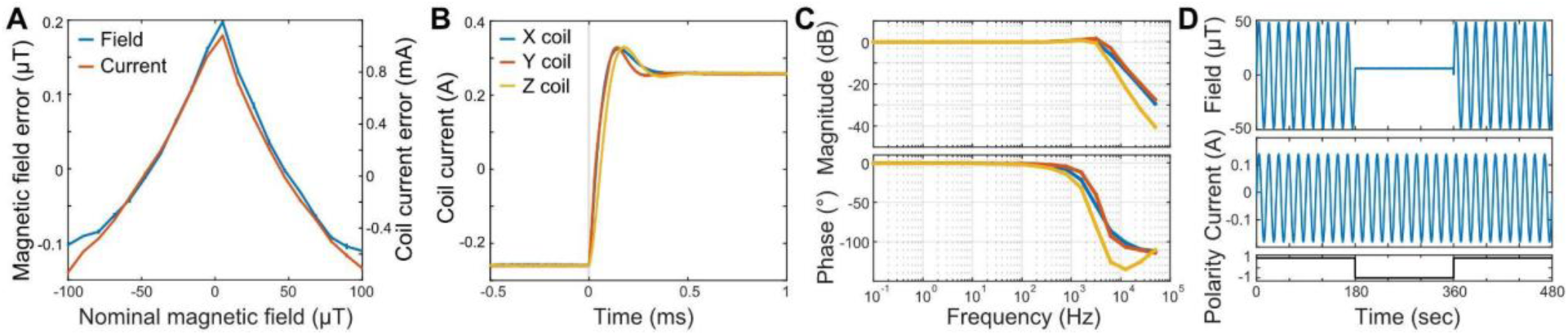
Coil amplifier. **A:** Coil amplifiers show an approximately linear response over a large range. Magnetic field strength and coil current were measured for a control voltage sweep corresponding to a magnetic field strength of approx -100 µT to 100 µT. Residuals of a linear fit of magnetic field and coil current are shown. Average of 3 repeats, error bars: ±SD. **B:** Coil current responses to a control voltage step for the three coils corresponding to a transition from -100 to 100 µT. Frequency compensation was adjusted for maximal speed of the transition and minimal overshot. **C:** Bode plots of amplitude attenuation (top) and phase shift (bottom) of the coil current for sinusoidal control voltages from 0.1 Hz to 50 kHz at ± earth magnetic field strength. **D:** Field blanking. By switching the coil wiring from serial to anti-serial (bottom) the magnetic field produced by the coils is canceled out (top) though the coils are still under current (middle). Note that the offset corresponds to the remaining natural magnetic field.

For the exemplary 3-axes coil system presented here, we adjusted the frequency compensation (see methods) to allow a moderate overshoot of ∼15% on all axes, bringing the system in a steady state below 500 µs (Fig. 3B). We applied sinusoidal control voltage oscillations between 0.1 Hz and 50 kHz with an amplitude corresponding to a steady state magnetic field strength of ∼50 µT and measured the amplitude and phase shift of the resulting sinusoidal coil current (Fig. 3C). In accordance with the step response, the frequency response showed only minor amplitude attenuation and phase shift up to 1 kHz.

By means of a coil polarity reversal relay, the magnetic field produced by the Helmholtz coils can be blanked while the windings are still under current. This is commonly used as an experimental control condition (sham magnetic stimulus, e.g. 36–39). If activated, the magnetic field is reduced to the ambient field strength present inside the setup (Fig 3D).

### Setup calibration

Control voltage vectors, which were uniformly distributed on a sphere, were applied to the coil drivers to calibrate the coil system. The resulting magnetic vectors inside the setup were measured at the location of the specimen with the miniature vector magnetometer. Similar to the magnetometer calibration, they resembled a distorted sphere, i.e., an ellipsoid, due to differing coil gains for the three axes and soft- and hard-iron distortions by components inside the setup. The parameters describing the transformation from the control voltages to the effective magnetic field strength at the position of the specimen were determined by linear regression. These parameters were then used to derive the coil amplifier control voltages that resulted in the desired magnetic vectors (see methods). The magnetic target vectors with a magnitude of 50 µT each were compared to the resulting vectors (Fig. 4A). The field error between target and actual vectors in magnitude was 0.003 ± 0.142 µT (Fig. 4B) and 0.191 ± 0.09 µT (mean ± std) in Euclidean distance (Fig. 4C).

**Figure 4:**
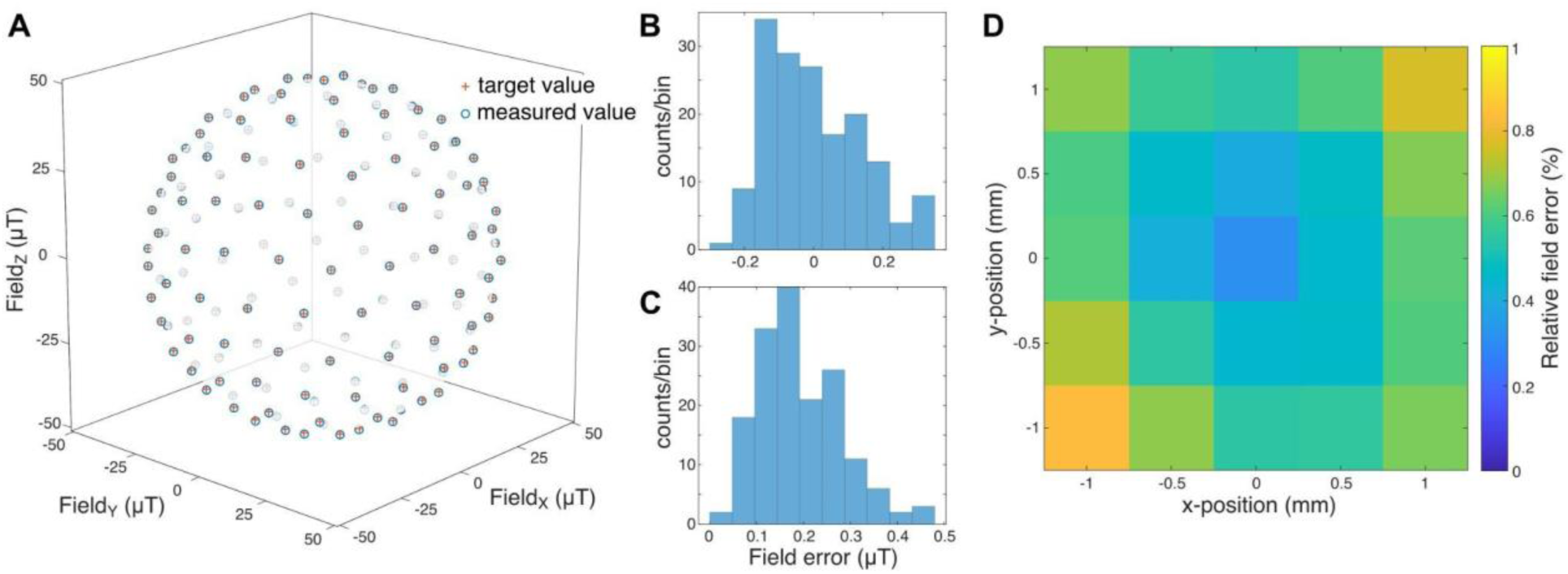
Setup calibration. **A:** Precision of 162 uniformly distributed field vectors in the calibrated setup. Red crosses: target vector, blue circles: measured field vector **B:** Magnitude deviations from the target of 50 µT. **C:** Euclidean distance between target and measured field vector. **D:** Field homogeneity in a 2 mm * 2 mm area in the recording chamber. At each location a spherical magnetic field stimulus with 44 vectors was analyzed. For each location, the mean Euclidean error is shown in percent of the target radius.

To test the spatial homogeneity of the generated field we consecutively placed the magnetometer in a grid of 5*5 positions in increments of 500 µm. We measured the accuracy of the stimulation at those locations and the average field error was determined (Fig. 4D). For the sampled area, the field error was below 1%.

To demonstrate the potential of the presented calibration technique, we purposefully introduced a strong source of magnetic field distortion into our physiological setup and corrected its field-deteriorating effects as far as possible (Fig. 5). We placed a 100 mm × 25 mm × 25 mm (approx. 500 g) ferromagnetic steel (mild steel) bar in two orientations (parallel to the magnetic y-axis, and parallel to the z-axis) in close proximity to the recording chamber inside our setup, which was beforehand calibrated without the metal bar, and estimated the accuracy as before. As expected, the metal bar led to strong distortions of the stimulus in both positions. By recalibrating the coil system in presence of the steel bar, the mean Euclidean error could be reduced to 0.5% (distortion 1) and to 0.6% (distortion 2), respectively. However, the spatial inhomogeneity, measured as in figure 4D over an area of 2 mm × 2 mm, was increased to 5%.

**Figure 5:**
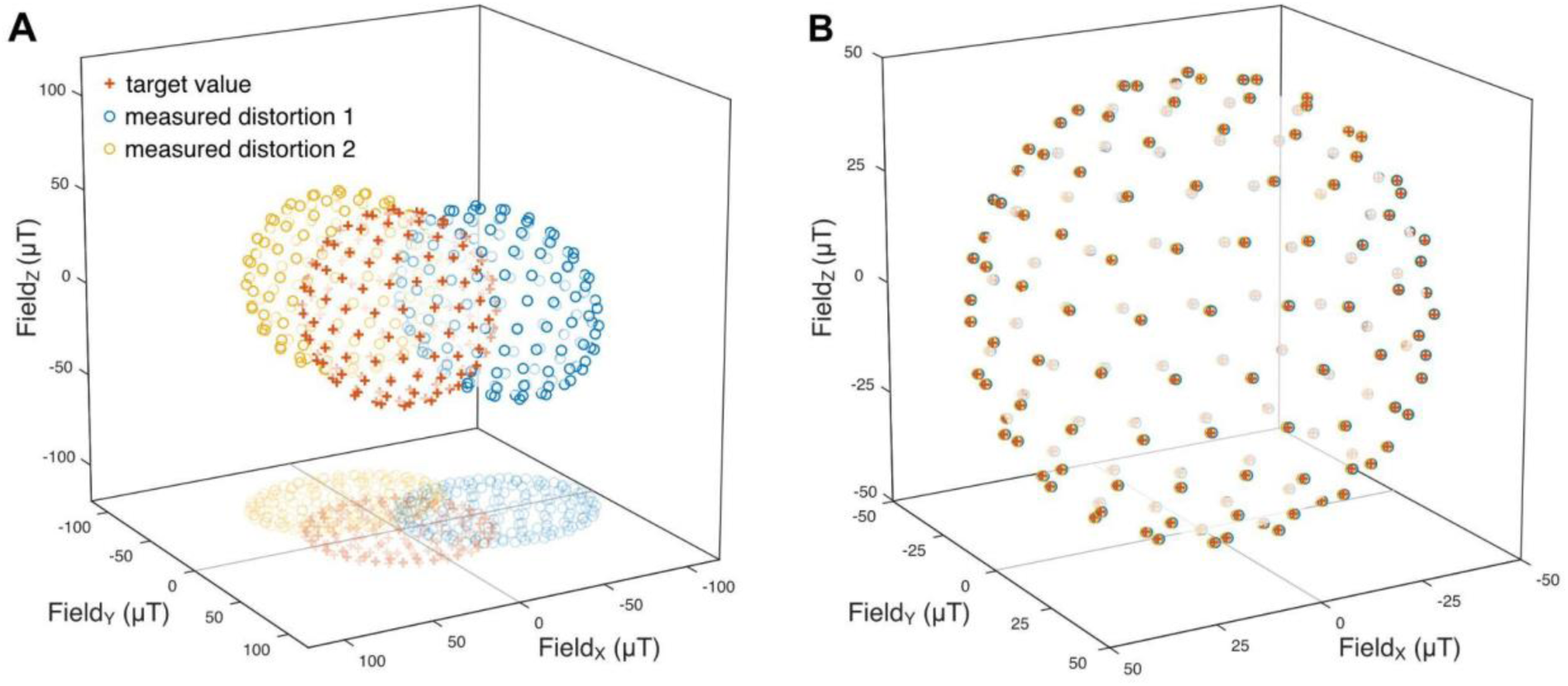
Compensation of strong distortions. **A:** A ferromagnetic steel bar was placed inside a previously calibrated setup in two orientations and a magnetic sphere stimulus with a target radius of 50 µT was applied (red crosses). The resulting sphere was offset and deformed for both positions of the metal bar (blue and yellow circles). **B:** After calibration, offset and deformation of the sphere were largely reduced, yielding average errors of 0.63% (distortion 1, blue circles) and 0.51% (distortion 2, yellow circles).

### Induction artifacts

Finally, with the generation and verification of the magnetic stimulation under control, the use of magnetic stimulation in electrophysiological methods faces another obstacle. Magnetic field transitions induce currents in conductive material, and the high impedance recordings common in electrophysiology are particularly sensitive to these small inductive artifacts. They might be picked up by the electrodes themselves, the headstage, or connecting cables. Therefore, one has to separate potential biological responses to the magnetic field transitions from the induced artifacts. The most straightforward difference between these two is the small latency between the field transition and the induction artifact in comparison to latencies of biological origin. However, this can only be exploited in certain experimental designs. Another distinctive feature may be the temporal waveform, as one would assume that induction artifacts have very different temporal waveforms in comparison to biological responses. In contrast, due to the low pass filter properties of the recording equipment, these waveforms can indeed be quite similar when responses on individual electrodes are considered. However, if one studies the signals simultaneously on multiple electrodes, the difference becomes quite apparent in all cases (Fig. 6A-D). Biological spikes originate at the axon hillock, backpropagate through the dendritic tree, and travel along the axon. The same spike is recorded differently on multiple electrodes while the induction artifact is observed over large areas identically and simultaneously. Common spike sorting algorithms that include simultaneous signals from multiple electrodes as from tetrodes or multi-electrode arrays generally classify artifacts and spikes reliably, as the signals are in this regard quite different (Fig. 6E). The strength of the induction strongly depends on the particular arrangement of the recording equipment relative to the coils. In our case, the strongest induction was seen for strong transitions in the coils in both X and Y direction, while the coils for the Z direction had little influence (Fig. 6D) ([x, y, z] = [-0.74, 0.67, 0.07], R^2^ = 0.990). This coincides with the planar structure of the electrode array and the PCB of the headstage. As expected, activation in the opposite polarity of the X and Y coils lead to an induction of opposite polarity.

**Figure 6:**
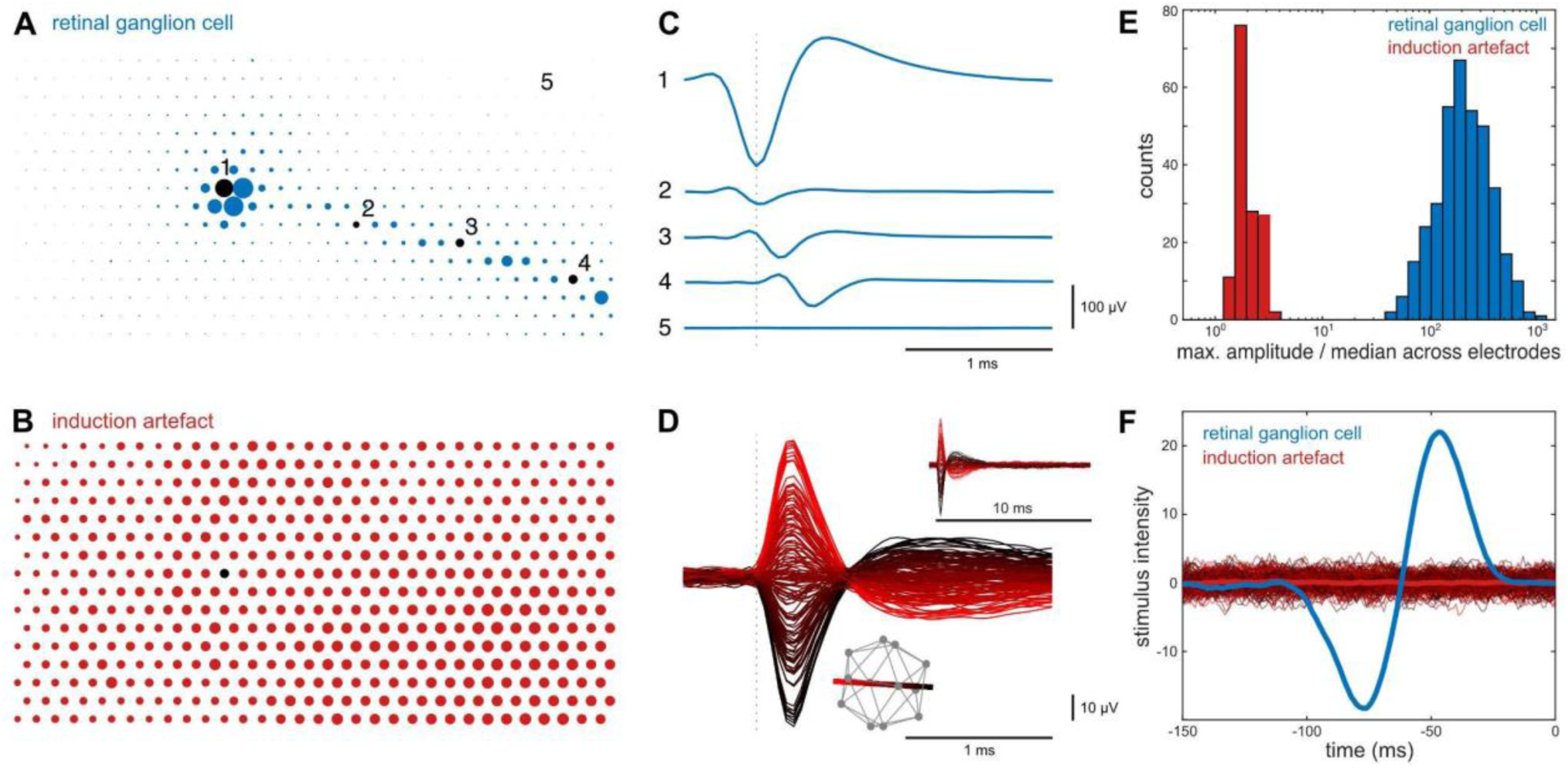
Rapid transitions of the magnetic field induce distinct electrical artifacts in the recording equipment. Multi-electrode array recording of quail retinal ganglion cells. **A:** Representation of the signal strength of a ganglion cell spike across the 512 electrodes of a multielectrode array. Dot size represents absolute signal amplitude. For visual clarity, maximal amplitudes are shown smaller than to scale. **B**: As A for an induction artifact. The black dot corresponds to position 1 in A. **C:** Average spike waveform on five electrodes marked in A. Note the strong signal at the cell soma location (1) and the increasing delay of the signal on the electrodes along the axon (2-4). **D:** Temporal induction artifact on electrode 1 as in C for 12*12 magnetic field transitions. Dashed line indicates the time of field transition. Bottom inset illustrates the icosahedron of the 12 used magnetic field vectors. The red-to-black line represents the axis with the strongest inductive effect and is used as the color map. Top inset: The same induction artifact at a longer time scale. **E:** Biological spikes (blue, n=367 quail retinal ganglion cells) and induction artifacts (red, n=144) have different spatiotemporal waveforms. Note logarithmic binning. **F:** Event-triggered average visual stimulus for spikes of the neuron in A (blue) and all magnetic field transitions (color map as in D). Thick red line: mean over all field transitions. Triggered average stimulus confirmed a clean classification with a normal light ON response for the neuron in A and no correlation between the visual stimulus and the induction artifacts, as expected.

## Discussion

We presented a system for magnetic stimulation and stimulus verification for physiological experiments, consisting of a miniature vector magnetometer and a current driver for electromagnetic coils, e.g., Helmholtz coils.

Due to tight space constraints in physiology research setups, non-ideal placement and/or geometry of the magnetic field generating coils is often necessary. Furthermore, the presence of field-disturbing ferromagnetic components inside the setup can be rarely avoided completely. Thus, it is important to verify the magnetic stimulus with a miniature magnetometer probe at the position of the specimen. Three-axes magnetometers based on small-sized Hall-effect sensors exist but are in general significantly less sensitive than e.g. fluxgate magnetometers. The latter are thus typically used for measuring small fields but are in turn bulkier. Therefore, we use a small-sized AMR three-axis magnetic field sensor to build a vector magnetometer whose sensor unit is small enough to fit in typical recording chambers of physiological setups, while being sufficiently sensitive to measure small fields. The sensory structures inside the sensor package are below 1 mm, resulting in a correspondingly small spatial integration area of the magnetic measurements. This is beneficial for the analysis of stimuli for small size biological specimens since it offers higher spatial resolution than larger magnetic probes. Only if ferromagnetic components that are smaller than the magnetometer’s spatial integration area are brought in close proximity to the specimen during stimulation, field inhomogeneities on scales below the spatial resolution of the sensor are to be expected. While purely analog AMR sensors were used in research on magnetoreception (16), the application of modern sensors with integrated analog frontend, A/D conversion, and digital data output, as presented here, greatly reduces the technological effort and thus facilitates the establishment of a robustly working system.

Optimally, the presented magnetometer is calibrated in a reference magnetic field provided by a calibrated coil system. Since such a system might not be readily available in physiology labs, we adapted a simple calibration method that is commonly used for smartphone integrated magnetometers. It has the advantage that it can easily be performed in a homogenous field of known magnitude, like the Earth’s magnetic field.

The used QMC5883L sensor was originally intended as a magnetometer for high-volume consumer applications like smartphones with power consumption and space constraints. While its small size is a key feature for the context described here, it cannot be expected to achieve the accuracy, bandwidth, and noise properties as dedicated devices like commercial fluxgate magnetometers. However, the typical physiology laboratory-use scenario of the magnetometer as a stimulus calibration and verification instrument allows long data averaging periods or repeated measurements. By this means, we achieved an accuracy of approx. 0.5% of the natural magnetic field.

We introduce a bipolar coil current driver design that is able to power coils with up to ±3A. The current range is optimized for small to medium-sized coil systems typically used in physiology setups, avoiding over-dimensioned control and supply devices for this use case. Contrary to the at times employed approach of constant voltage supply of coil systems, our constant current driver circuit eliminates the problem of decreasing magnetic output caused by coil heating through ohmic losses during operation. As a commonly used control for one type of possible artifacts, the driver offers polarity reversal of one winding of a double-wrapped coil, thereby blanking the magnetic field generation while keeping the coils electrically active. Due to its modular approach, the coil driver circuit can easily be integrated into running systems, if free analog-out channels for field control are available. Any standard bipolar lab power supply unit can be used to power the driver if the required voltages and currents can be delivered. The circuitry can simply be multiplied according to the number of magnetic stimulation axes desired, hence systems with 1-, 2-, or 3-dimensional stimulation can flexibly be set up. We are successfully using the amplifier design for three-axis quadratic Helmholtz-type magnetic stimulation in a multi-electrode electrophysiology setup. While coil system designs according to Lee-Whiting, Merrit, Alldred and Scollar, or Rubens offer better field homogeneity over larger areas than Helmholtz coils, they require more coils (between 3 and 5) per axis (20). Thus, for space-constricted *in vitro* physiology setups, Helmholtz coil systems have the advantage of occupying less space than the other designs. The disadvantage of a smaller region of high field homogeneity is typically acceptable since the specimens of *in vitro* physiology are small in most cases. While we applied the presented coil driver to a square Helmholtz-type coil arrangement, the driver can be used with other coil arrangements within the current specifications.

In general, heat production by the coil system is a concern in bioelectromagnetic studies (40). Since temperature is an omnipotent physiological parameter (41), temperature effects on the specimen caused by coil heating may obscure real effects or be mistaken for them. However, this is an issue primarily in studies with field strengths significantly higher than the earth’s magnetic field. In case of our exemplary Helmholtz coil system, no significant temperature increase of the coils was apparent in their normal operational range. Furthermore, the coils are not in heat-conducting contact with the recording chamber, and the specimen is often temperature controlled in physiology setups. For certain stimulus conditions one can exploit the fact that a sham stimulation using double-wrapped coils produces the same amount of heat as the non-sham stimulation.

If disturbances of the generated magnetic stimulus are identified by the magnetometer, a simple setup calibration technique can be applied which compensates for stationary soft- and hard-iron disturbances. We applied the calibration technique to our multi-electrode electrophysiology setup as an example of a typical physiology research setup. While we avoided the use of ferromagnetic materials when building the setup, small amounts are present in the used microscope objectives and in the recording electronics in close proximity to the preparation. After setup calibration, we achieved an average field error below 0.4% of a magnetic vector of a given magnitude and a spatial field homogeneity better than 1% in an area of 2 mm * 2 mm at the position of the recorded specimen. This is more than sufficient for the physiological study of magnetoreception in the foreseeable future. These results obviously depend on the properties of the specific setup. In addition, instead of calibrating the field for one central region, one could alternatively optimize homogeneity of the full recording area at once. To further elucidate the potential severity of distortions inside the setup, we compensated for extreme field disturbances introduced by a ferromagnetic steel bar of approx. 500 g. In this case, the magnitude error for a given location was still below 1%. But as expected, spatial field homogeneity was more strongly compromised (5% error in an area of 2 mm * 2 mm). While these strong distortion sources are obviously an exaggeration of what might realistically be present in an actual research setup, it shows that even these can be compensated to a potentially acceptable degree by the suggested calibration technique. In fact, the local stimulus quality is within the acceptable limits for experiments on smaller specimens, like single-cell recordings.

Since magnetic artifacts in recordings are induced by magnetic field changes, fast transition times between magnetic stimuli help shorten the artifacts. Hence, if instantaneous magnetic stimulus changes are applied in a recording, they will occur transiently with very short latencies in comparison to biological responses and can be excluded from the analyses. On the other hand, if slow magnetic stimuli are applied, the induced artifacts can be expected to be smaller in amplitude because less change of the magnetic field per time will result in smaller artifact induction (42). Magnetic stimulus transitions might be chosen to be slow enough to fall below the physiological recording’s frequency band. Thanks to its wide frequency band our coil driver is well suited for both approaches. If possible, the induction artifact should be further minimized e.g. by the arrangement of the cables or the use of twisted-pair cables (42). If spike sorting with multiple electrodes is possible, the induction artifact is easy to separate from biological signals. However, due to the filter properties of the stimulation and recording devices, the artifact waveform on a single electrode can resemble a biological spike, and particular care in the experimental design is needed. Additionally, control conditions like the pharmacological block of synaptic transmission or cooling of the neuronal tissue can be applied to distinguish between stimulation artifacts and physiological responses.

In combination with commonly available computer-controlled digital-to-analog interfaces, our coil driver allows easy implementation of complex stimulation paradigms that can be applied automatized to the studied specimen. In our exemplary multi-electrode recording setup, we simultaneously stimulate the specimen visually and magnetically. By this means, our setup enables the generation of synchronized multimodal stimuli that, for example, can simulate natural animal behaviors like birds’ head scanning movements before take-off (43). On the other hand, due to the fast response time of the coil amplifier, also less natural system-theory-inspired stimulation paradigms can easily be implemented, like randomly fast changing magnetic noise stimuli for reverse correlation analyses. While our coil driver design was dimensioned according to the demands of physiological *in vitro* research setups, it can as well be applied to *in vivo* experimentation.

Magnetic stimulation is error-prone since the stimuli are not directly detectable by the experimenter, as opposed to visual stimuli, for instance. Therefore, it is good practice to monitor the magnetic field during the experimental procedure to avoid stimulation errors (16). Due to the digital data transmission between the sensor PCB and the read-out electronics of the magnetometer (Fig. 1B), the connecting cable can be several meters long without impairing the quality of the measured data. In addition, the small size makes the magnetometer well suited to be permanently installed inside a physiology setup for stimulus control. If required by the experimental design, a closed-loop stimulation can be implemented with minimal effort. If magnetometer read-out and stimulation control are performed by the same software platform (e.g. Matlab), the magnetometer data can easily be fed back into the stimulus generation. This is beneficial if the exact placement of setup components cannot be kept unaltered over experiments, which would otherwise require a recalibration of the system.

However, in this case, the overall timing constraints of the magnetometer’s data transmission (maximum data output rate of 200 Hz, USB latency, and jitter) have to be evaluated in regard to the intended stimulation.

While the presented methodology was developed in the context of multi-electrode recordings, it can easily be used with other physiological techniques that share similar inherent problems in regard to the generation and evaluation of magnetic stimuli. In conjunction, the presented miniature vector magnetometer, coil driver, and coil calibration technique are well suited to establish a reliable magnetic stimulation system for a wide variety of physiology setups, i.e. for strongly space-constricted setups dedicated to experiments with small specimens.

## Acknowledgements

We thank C. Puller and H. Mouritsen for their valuable comments on the manuscript. Research was supported by SFB 1372, Deutsche Forschungsgemeinschaft (MG and MW) and RTG 1885/2, Deutsche Forschungsgemeinschaft (MG and MW).

The complete hardware design data and software resources are open source under the Creative Commons Zero v1.0 Universal license (www.github.com/mtahlers/magStim).

